# *In-situ* genomic prediction using low-coverage Nanopore sequencing

**DOI:** 10.1101/2021.07.16.452615

**Authors:** Harrison J. Lamb, Ben J. Hayes, Imtiaz A. S. Randhawa, Loan T. Nguyen, Elizabeth M. Ross

**Affiliations:** Centre for Animal Science, Queensland Alliance for Agriculture and Food Innovation, The University of Queensland, Brisbane, QLD 4072, Australia; School of Veterinary Science, The University of Queensland, QLD, 4343, Australia

## Abstract

Most traits in livestock, crops and humans are polygenic, that is, a large number of loci contribute to genetic variation. Effects at these loci lie along a continuum ranging from common low-effect to rare high-effect variants that cumulatively contribute to the overall phenotype. Statistical methods to calculate the effect of these loci have been developed and can be used to predict phenotypes in new individuals. In agriculture, these methods are used to select superior individuals using genomic breeding values; in humans these methods are used to quantitatively measure an individual’s disease risk, termed polygenic risk scores. Both fields typically use SNP array genotypes for the analysis. Recently, genotyping-by-sequencing has become popular, due to lower cost and greater genome coverage (including structural variants). Oxford Nanopore Technologies’ (ONT) portable sequencers have the potential to combine the benefits genotyping-by-sequencing with portability and decreased turn-around time. This introduces the potential for in-house clinical genetic disease risk screening in humans or calculating genomic breeding values on-farm in agriculture. Here we demonstrate the potential of the later by calculating genomic breeding values for four traits in cattle using low-coverage ONT sequence data and comparing these breeding values to breeding values calculated from SNP arrays. At sequencing coverages between 2X and 4X the correlation between ONT breeding values and SNP array-based breeding values was > 0.92 when imputation was used and > 0.88 when no imputation was used. With an average sequencing coverage of 0.5x the correlation between the two methods was between 0.85 and 0.92 using imputation, depending on the trait. This demonstrates that ONT sequencing has great potential for in clinic or on-farm genomic prediction.

**Author Summary:** Genomic prediction is a method that uses a large number of genetic markers to predict complex phenotypes in livestock, crops and humans. Currently the techniques we use to determine genotypes requires complex equipment which can only be used in laboratories. However, Oxford Nanopore Technologies’ have released a portable DNA sequencer, which can genotype a range of organisms in the field. As a result of the device’s higher error rate, it has largely only been considered for specific applications, such as characterising large mutations. Here we demonstrated that despite the devices error rate, accurate genomic prediction is also possible using this portable device. The ability to accurately predict complex phenotypes such as the predisposition to schizophrenia in humans or lifetime fertility in livestock *in-situ* would decrease the turnaround time and ultimately increase the utility of this method in the human clinical and on-farm settings.

## Introduction

Complex traits in livestock, crop and human genetics are primarily polygenic, that is, a large number of variants contribute to genetic variation. These traits, which are often continuous (e.g., height [1], weight [2] or temperament [3]) are heritable to varying degrees [4]. The contribution to the overall phenotype of each variant typically ranges between very small effects for common variants to larger effects for some low frequency variants [5]. Each allele an individual carries at loci associated with the complex trait contributes to an increase or decrease in the phenotype [4]. By exploiting this relationship, it is possible to predict the complex phenotype of a genotyped individual without a phenotype if the variant effects are estimated using appropriate statistical methods such as genomic best linear unbiased prediction, Bayesian methods, or polygenic risk scores in humans [6, 7].

In livestock and crops, genomic predictions are termed genomic estimated breeding values (GEBVs) and are used to increase the accuracy and intensity of selection in a process referred to as genomic selection [6]. The benefits of genomic prediction include more accurate selection of complex traits and a decrease in generation interval, which ultimately leads to accelerated genetic gain, evidenced in the poultry [8] and dairy industries [9]. The method allows for the polygenic nature of complex traits by genotyping a number of markers spread evenly throughout the genome and assigning an effect to each marker.

Today, almost all livestock species have SNP arrays designed to enable genomic selection. In Australia, the turnaround time between sample collection and GEBV results is between 6-8 weeks. This turnaround time has prevented the adoption of genomic selection in Australia’s northern beef industry, where cattle are generally only handled once a year for a handful of days. This means management decisions based on the GEBV results from SNP array genotyping cannot be implemented until the following year, when the cattle are handled again. There is also significant demand for faster GEBV turnaround time in the sheep and southern beef industries to allow point of management decisions, for example, allocation of feeding regimes based on genetic potential.

In human genetics, complex disease traits have been the focus of this type of quantitative genetics, although pharmacogenetics also holds some potential. For complex diseases such as bipolar disorder, Type II Diabetes, and Crohn’s disease [10] polygenic risk scores are used to evaluate an individual’s genetic predisposition to the disease. Poly-genic risk scores are based on the genome wide association study (GWAS) correlation between significant SNP markers and the disease [11]. To-date the large-scale utility of polygenic risk scores for individuals has been limited [5]. This has been attributed in part to the current turnaround time, and cost. Instead, family history is used to estimate an individual’s predisposition to a particular disease.

However, family history information has limited utility for determining relative risk for complex diseases in some cases. For example, in schizophrenia a family history of the disease is reported in less than a third of cases [12]. A Swedish national study [13] reported family history of the disease in only 3.81% of cases, taking into account first-, second- and third-degree relatives. Moreover, despite half the genetic variance for schizophrenia occurring within family, siblings with a family history of the disease are given the same risk using family history alone. By genotyping the individual, polygenic risk scores far more accurately report an individual’s genetic risk than family history alone. In the case of schizophrenia, this information could be used to differentiate between individuals vulnerable to environmental risk factors [14].

Studies exploring the polygenic nature of pharmacological response to chemotherapeutics [15] and asthma treatments [16] have also yielded promising results. Decreasing the turnaround time of genotyping could increase the utility of polygenic risk scores for individuals. For example, a clinician may like to know the genetic risk of a particular disease to assess the priority of diagnostic tests, predict the efficacy of a potential drug treatment, or predict the patient’s likelihood of having a severe adverse reaction to a particular treatment, as quickly as possible before the treatment is due to be administered. Decreased turnaround time for poly genic risk scores would also help clinicians diagnose complex diseases in patients while in their prodromal phase, which is vital in early intervention [14].

Genomic prediction in both medicine and agriculture has previously relied heavily on SNP array technology. SNP arrays are a low-cost method to genotype thousands to hundreds-of-thousands of genetic markers. Once costing $400 USD for 10,000 SNP genotypes, SNP arrays now cost less than $50 USD for 50,000-800,000 SNP genotypes. However, the technology requires large, expensive laboratory equipment. Therefore, the turnaround time is, at a very optimistic minimum, the time for a sample to reach the laboratory.

Recently, there has been a rise in the popularity of genotyping-by-sequencing. A widely applied approach is to use restriction enzymes to reduce the complexity of genomes for low-coverage sequencing on short-read sequencing platforms [17]. This method is not only becoming cheaper than SNP array genotyping, but it also allows for simultaneous genome wide marker discovery. These advantages have allowed an increased number of traits and associated variants to be studied [4], which has resulted in the ability to accurately predict many new complex traits from genotypes. Still, genotyping-by-sequencing is laboratory based and requires expensive equipment. A potential solution to reduce the turnaround time of genomic prediction while incorporating the benefits of genotyping-by-sequencing, is to use Oxford Nanopore Technologies’ (ONT) portable nucleotide sequencer, the MinION [18].

The MinION could be used to sequence samples *in-situ* for genomic prediction in livestock and human settings. Despite initial reports of poor sequencing accuracy and yield, steady developments in flow cell chemistry, library preparation and base calling algorithms has seen reported sequencing yields increase from less than 3 GB to greater than 40 GB and sequencing accuracy increase from 68.4% [19] to over 98% [20, 21]. A major advantage of ONT sequencing technology is the ability for ONT reads to map more accurately to complex genomic regions [22]. This is largely a result of the read length produced by the technology, which has no theoretical limit. This reduces read mapping bias [23] and eliminates the need for restriction enzyme digestion used for genotyping-by-sequencing with short read sequencing technology.

The aim of this study was to evaluate the accuracy of genomic predictions calculated from low-coverage ONT sequence data and compare the results to SNP array-based predictions. We calculated correlations and prediction bias for ONT genomic predictions against the SNP array predictions for various sequencing depths (4x, 2x, 1x, 0.5x) and tested three different methods of imputing missing genotypes. Our results suggest ONT’s MinION sequencer could be a useful tool for *in-situ* genomic prediction, with human clinical or on-farm applications.

## Results

### Sequencing

ONT sequencing of 19 cattle from tail hair yielded an average read length of 1,797 bp and an average flow cell yield of 22.57 Gb over the 96-hr run, with the highest yield being 41.13 Gb (Table 1). The average Phred scaled base quality was 20.54 ± 0.16 with a maximum of 22.3 and minimum of 18.2. On average 86.4% ± 0.6 of reads were effective (i.e., mapping quality Phred score > 0). For each animal, a random subset of sequence data representing 4x, 2x, 1x, and 0.5x sequencing coverage of the 2.7 Gb bovine genome was used for genomic predictions.

**Table 1:**
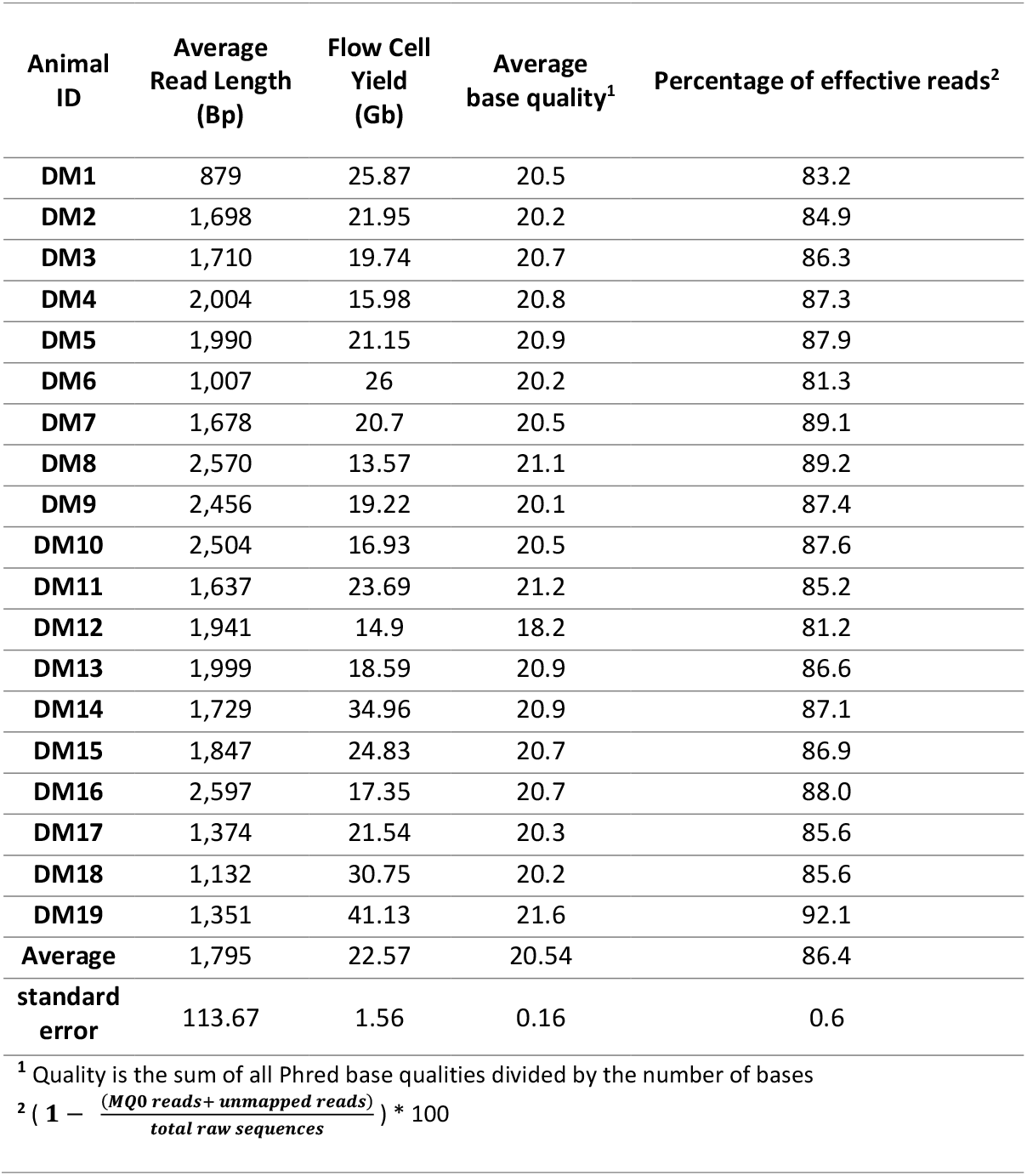
Read length and flow cell yield for each sample sequenced on the MinION.

| Animal ID | Average Read Length (Bp) | Flow Cell Yield (Gb) | Average base quality<sup>1</sup> | Percentage of effective reads<sup>2</sup> |
| --- | --- | --- | --- | --- |
| DM1 | 879 | 25.87 | 20.5 | 83.2 |
| DM2 | 1,698 | 21.95 | 20.2 | 84.9 |
| DM3 | 1,710 | 19.74 | 20.7 | 86.3 |
| DM4 | 2,004 | 15.98 | 20.8 | 87.3 |
| DM5 | 1,990 | 21.15 | 20.9 | 87.9 |
| DM6 | 1,007 | 26 | 20.2 | 81.3 |
| DM7 | 1,678 | 20.7 | 20.5 | 89.1 |
| DM8 | 2,570 | 13.57 | 21.1 | 89.2 |
| DM9 | 2,456 | 19.22 | 20.1 | 87.4 |
| DM10 | 2,504 | 16.93 | 20.5 | 87.6 |
| DM11 | 1,637 | 23.69 | 21.2 | 85.2 |
| DM12 | 1,941 | 14.9 | 18.2 | 81.2 |
| DM13 | 1,999 | 18.59 | 20.9 | 86.6 |
| DM14 | 1,729 | 34.96 | 20.9 | 87.1 |
| DM15 | 1,847 | 24.83 | 20.7 | 86.9 |
| DM16 | 2,597 | 17.35 | 20.7 | 88.0 |
| DM17 | 1,374 | 21.54 | 20.3 | 85.6 |
| DM18 | 1,132 | 30.75 | 20.2 | 85.6 |
| DM19 | 1,351 | 41.13 | 21.6 | 92.1 |
| Average | 1,795 | 22.57 | 20.54 | 86.4 |
| standard error | 113.67 | 1.56 | 0.16 | 0.6 |

<sup>1</sup> Quality is the sum of all Phred base qualities divided by the number of bases
<sup>2</sup> $(1 - \frac{(MQ0\ reads + unmapped\ reads)}{total\ raw\ sequences}) * 100$

A minimum allele observation based genotyping approach was used to genotype 641,163 SNP markers. This method accounted for the random probability of sampling alleles in a diploid species by grouping loci with similar coverage together and calling genotypes using a minimum allele count specific to each coverage group. Three methods (non-imputed, imputed-AF & imputed-Beagle) were tested to genotype loci with less than one overlapping ONT read (where only one read was observed all heterozygous loci were called as homozygous regardless). Although the first two methods are obviously not accurate, they are very fast which could be useful on farm or in a clinical setting. Before imputation, at 0.5x sequencing coverage 64% of ONT genotypes were correctly called, 34% were incorrect in one allele (i.e., called homozygous rather than heterozygous) and 0.02% were incorrect in both alleles (opposing homozygous; Figure 1). At 4x sequencing coverage 74% of calls were correctly called in both alleles, 24% were called incorrectly in one allele and 2% were called incorrectly in both alleles.

**Figure 1:**
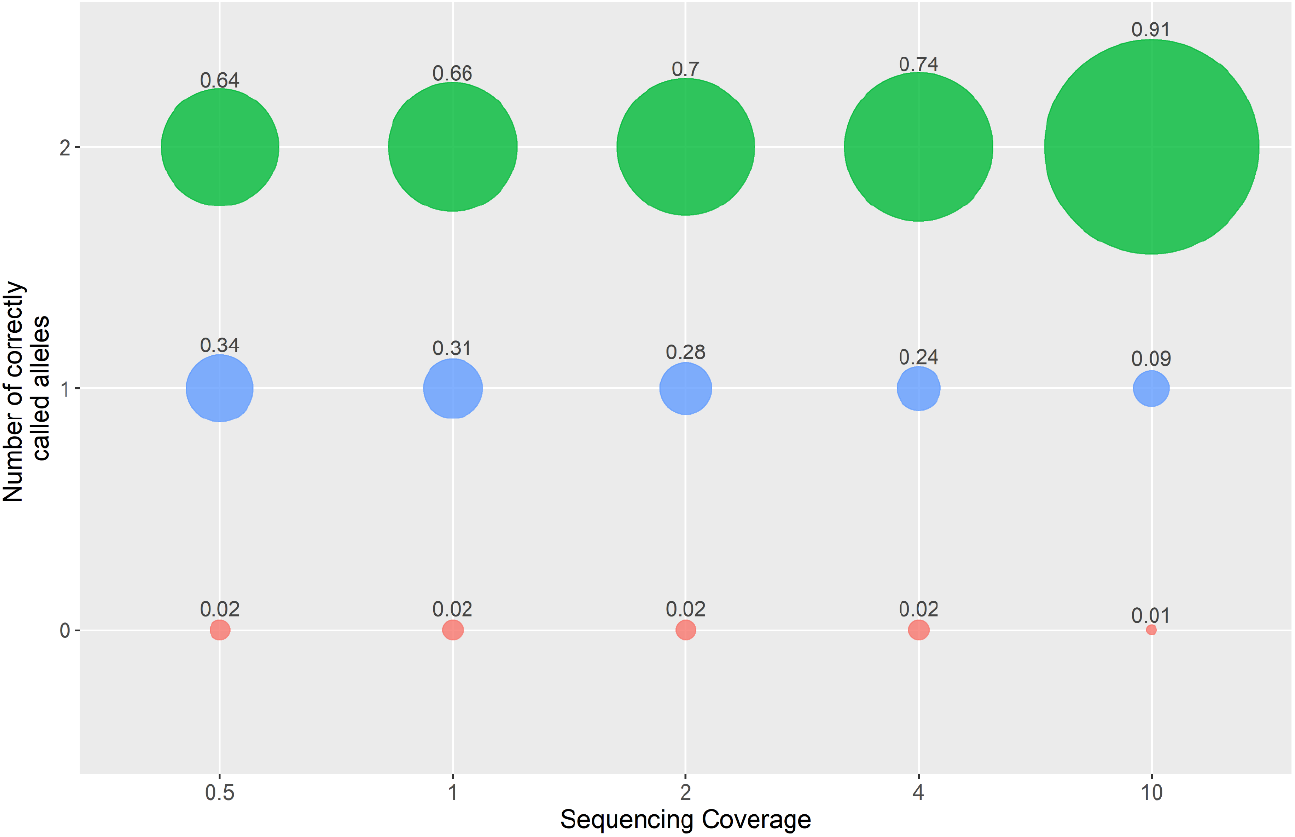
Proportion of genotype calls with both alleles correct (2), one allele incorrect (1) and two alleles incorrect (0) at different sequencing coverages. Two correctly called alleles (green) indicates no difference between the ONT genotype and SNP array genotype. One correctly called allele (blue) indicates one allele of the ONT genotype is correct and the other is incorrect. If both alleles are incorrect the (i.e., the alternate homozygous has been called) the number of correctly called alleles is zero (red).

### Marker Effects

Marker effects (BLUP solutions) for predicting GEBV were derived from 26,145 female cattle genotyped for 641,163 SNP markers from Hayes, Fordyce & Landmark [24] for each of the four traits: body condition score, body weight, corpus luteum at 600 days (CL600) and hip height. The marker effects were highly polygenic (Figure 2) with the largest marker effect for each trait less than 0.007% of the total effect. The hip height and body weight marker effects had the largest standard deviations: 0.0011 and 0.00063. While CL600 and body condition score had the smaller standard deviations: 0.00048 and 0.00041, respectively. These SNP marker effects were used to calculate genomic predictions for each of the four traits using ONT genotypes. The effect of read length, base quality, effective mapping percentage and number of reads mapping with quality 0 on ONT genomic prediction accuracy was tested. None of these covariates had a significant effect (linear model; P > 0.01) on the prediction accuracy.

**Figure 2:**
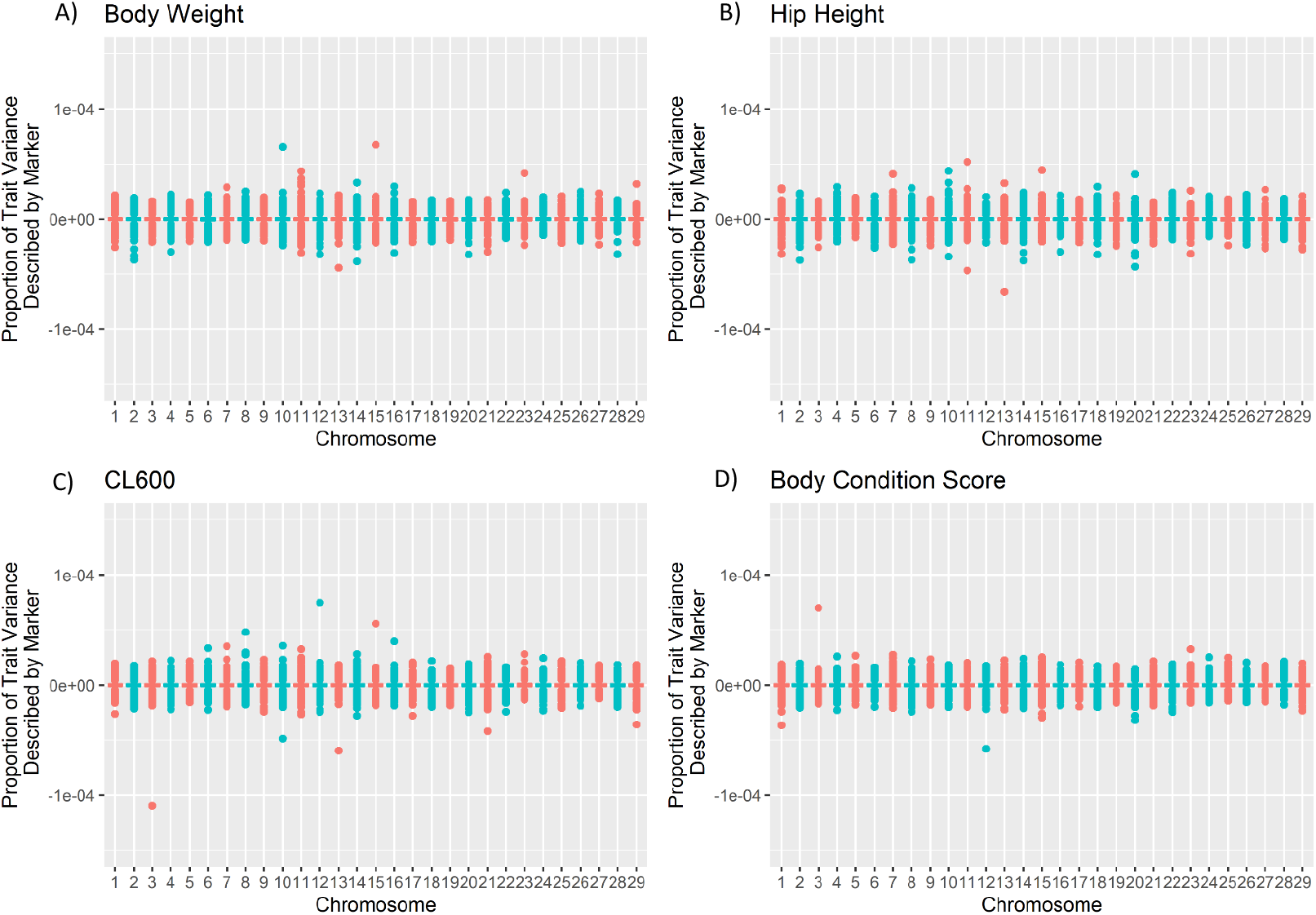
Proportion of trait variance described by the markers across the autosomes for each trait, where a positive value indicates a positive effect on the trait and a negative value a negative effect on the trait. The proportion of trait variance was calculated as 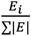 where *E_i_* is the effect of the *i_th_* marker and ∑|*E*| is the sum of the magnitudes of all markers for a particular trait. A) Body weight B) Hip Height C) CL600 D) Body Condition Score.

### Genomic prediction accuracies based on non-imputed genotypes

The non-imputed genotyping method, where uncalled loci were assigned a homozygous reference genotype, was as expected the least accurate for genomic prediction across all traits (Figure 3). The ONT non-imputed genomic predictions had reasonable correlations with the SNP array predictions from 4x down to 1x sequencing coverage (0.98 – 0.95 at 4x and 0.95 – 0.85 at 1x). These correlations dropped significantly at 0.5x sequencing coverage, 0.65 for body weight, 0.81 for CL600, 0.72 for body condition score and 0.69 for hip height (Figure 3). For sequencing coverages below 4x the prediction bias (regression coefficient of SNP array GEBV on ONT GEBV) for the non-imputed method was consistently greater than 1, indicating the ONT GEBV are under-estimating compared to the SNP array GEBV. At 2x sequencing coverage the bias ranged between 1.1 for body weight and 1.54 for body condition score (Figure 4; Table 2). At 0.5x sequencing coverage the bias for these two traits increased to 1.45 for body weight and 1.79 for body condition score. For the CL600 trait the bias at 0.5x was 2.7.

**Figure 3:**
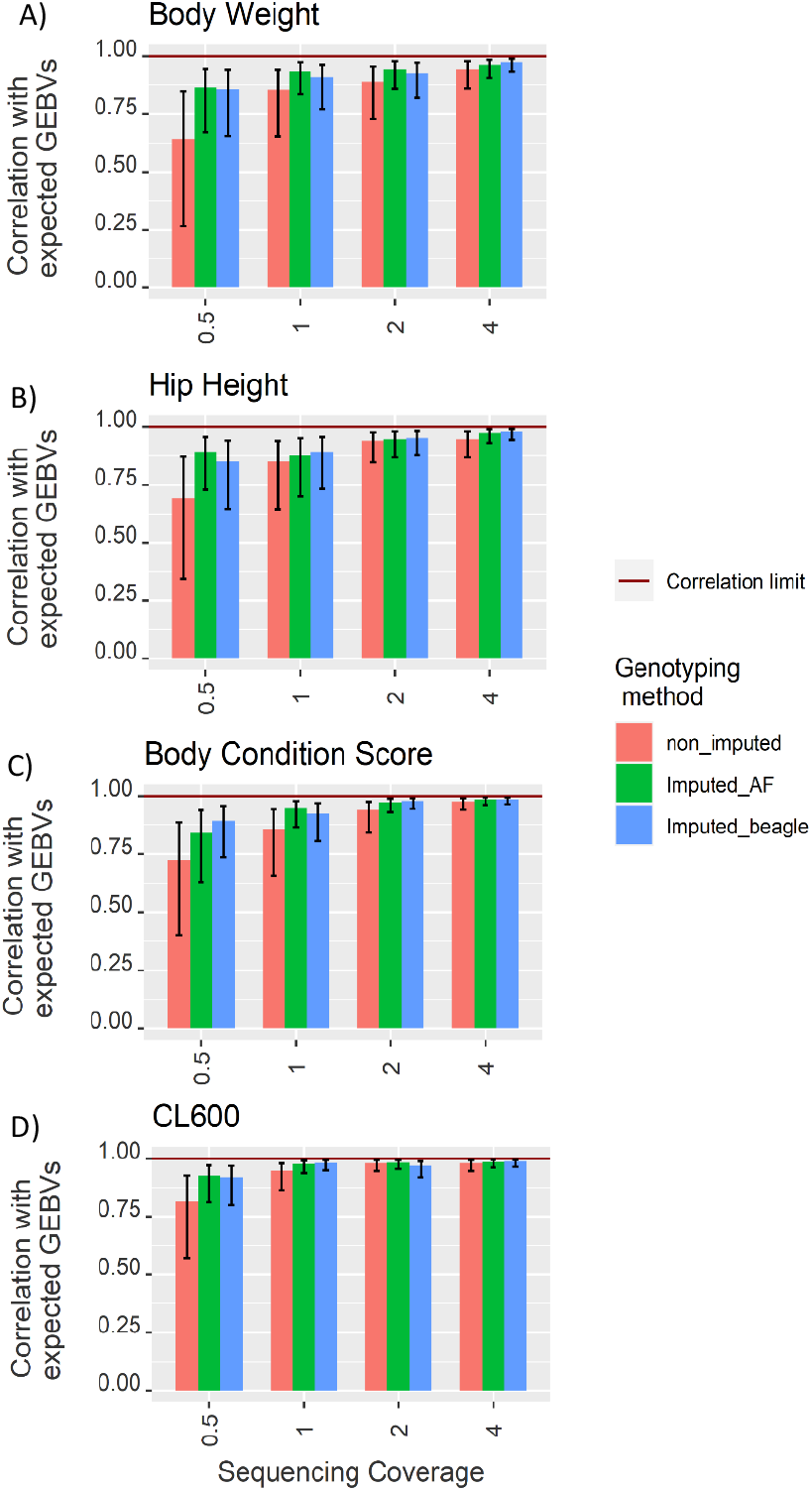
Correlations between the ONT genomic predictions and the expected GEBVs based on SNP array genomic predictions at four sequencing coverages for the four traits; A) body weight, B) hip height, C) body condition score, and D) CL600.

**Figure 4:**
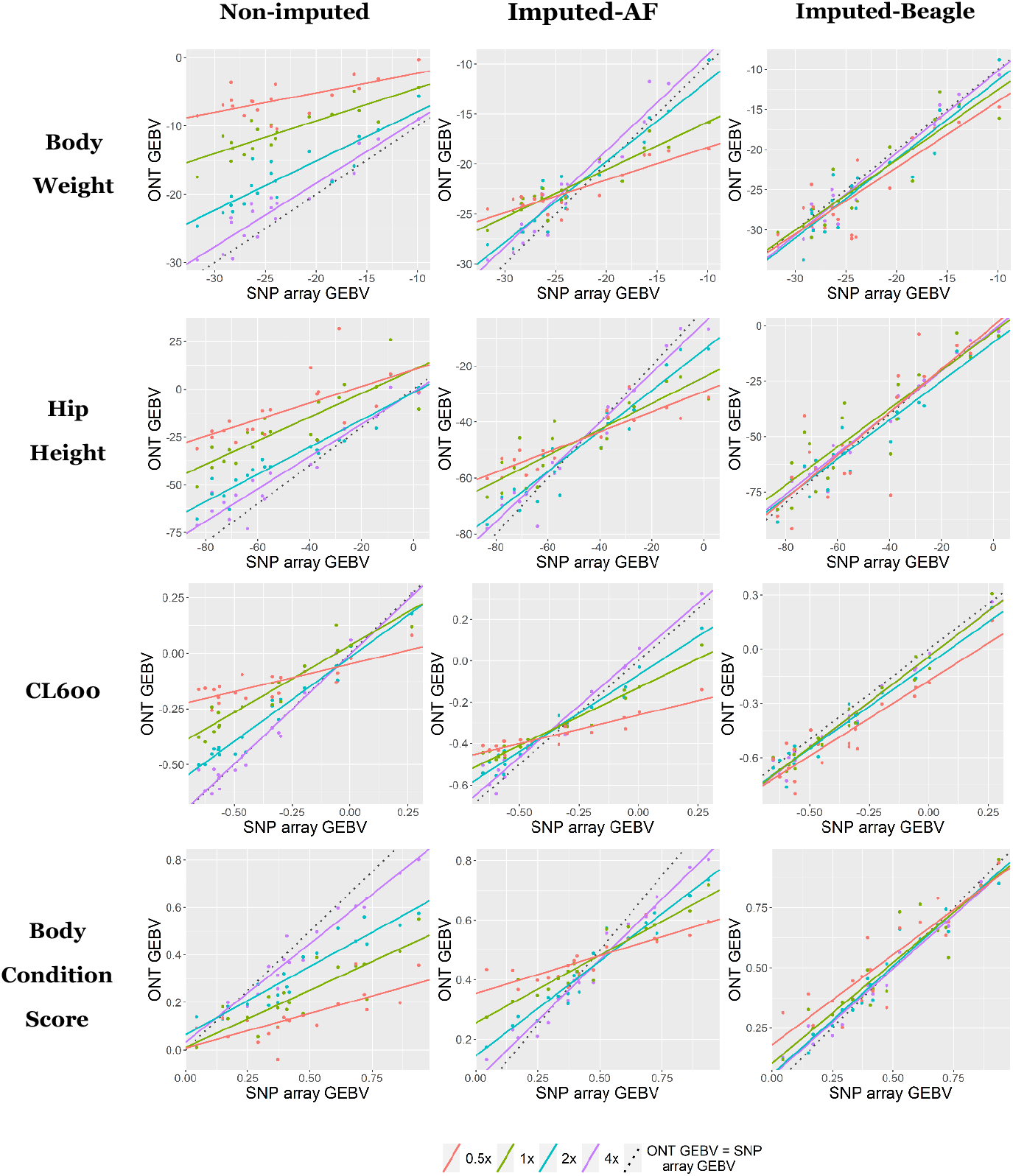
Correlations between the ONT genomic prediction and the SNP array genomic predictions for each of the four traits and three genotyping methods use.

**Table 2:** Prediction bias for the different genotyping methods and sequencing coverages across the four traits.

|  | Sequencing Coverage<br>(x of bovine genome) | Bias<br>(Regression coefficient of the SNP array-GEBV on the ONT-GEBV) |  |  |
| --- | --- | --- | --- | --- |
|  |  | Non-imputed | Imputed-AF | Imputed-Beagle |
| **Body weight** | 4 | 0.96 | 0.96 | 0.93 |
|  | 2 | 1.1 | 1.11 | 0.88 |
|  | 1 | 1.52 | 1.85 | 0.94 |
|  | 0.5 | 1.45 | 2.27 | 0.88 |
| **CL600** | 4 | 0.98 | 0.98 | 0.96 |
|  | 2 | 1.27 | 1.3 | 1.01 |
|  | 1 | 1.49 | 1.72 | 0.96 |
|  | 0.5 | 2.7 | 3.13 | 1.02 |
| **Body condition score** | 4 | 1.15 | 1.2 | 1.09 |
|  | 2 | 1.54 | 1.49 | 1.06 |
|  | 1 | 1.52 | 1.96 | 1.02 |
|  | 0.5 | 1.79 | 2.78 | 1.06 |
| **Hip height** | 4 | 1.06 | 1.08 | 1.03 |
|  | 2 | 1.23 | 1.23 | 1.03 |
|  | 1 | 1.16 | 1.64 | 0.92 |
|  | 0.5 | 1.1 | 2.22 | 0.74 |

### Genomic prediction accuracies based on imputed-AF genotypes

The correlation coefficients for the imputed-AF method, which used the population allele frequency to genotype missing loci, were the highest of the three methods. At 4x sequencing coverage the correlations were 0.99 for body condition score & CL600, 0.96 for body weight and 0.97 for hip height. This dropped slightly at 2x sequencing coverage, down to 0.95 for body weight, 0.98 for CL600, 0.97 for body condition score and 0.95 for hip height. The correlations remained above 0.9 for all traits at 1x sequencing coverage except for hip height, which was 0.88. At 0.5x sequencing coverage the correlations for this method remained above 0.8 with the lowest being 0.84 for body condition score. The prediction bias for this method was consistently greater than 1, ranging between 1.65 for hip height and 1.96 for body condition score at 1x sequencing coverage. This increased to between 2.2 and 3.1 for hip height and CL600 respectively at the lowest sequencing coverage.

### Genomic prediction accuracies based on imputed-Beagle genotypes

The final genotyping method, which used the imputation package Beagle v5.1 [25], had correlations greater than 0.85 for all traits at sequencing coverages as low as 0.5x. At 4x sequencing coverage the correlations for body weight, CL600, body condition score and hip height were 0.97, 0.99, 0.98 and 0.98 respectively. The correlations decreased slightly with the decrease in sequencing coverage to 0.85 for body weight, 0.91 for CL600, 0.89 for body condition score and 0.85 for hip height at 0.5x sequencing coverage. Although the correlations for this genotyping approach were, on average, not as high as the imputed-AF approach, the prediction bias was significantly reduced. The prediction bias at 4x sequencing coverage was around 1 for all traits and decreased slightly as the sequencing coverage decreased. The decrease in prediction bias was most notable in the body weight and hip height traits, which had a bias of 0.88 and 0.74 respectively at 0.5x sequencing coverage. For the same sequencing coverage CL600 and body condition score has a bias of 1.02 and 1.06.

## Discussion

In this study we sequenced 19 Droughtmaster heifers on ONT’s portable MinION sequencer, in order to compare genomic predictions from ONT data to SNP array genotyping, the current standard for genomic prediction. We investigated the accuracy of the ONT genomic predictions at various sequencing coverages (4x, 2x, 1x, and 0.5x) as well as using three different methods to genotype loci with no sequencing coverage: non-imputed, imputed-AF and imputed-Beagle. Beagle [25] imputation produced highly correlated GEBVs across all sequencing coverages. At 0.5x sequencing coverage this method had GEBV correlations greater than 0.85 and there was no evidence of prediction bias. On the other hand, the non-imputed and imputed-AF methods demonstrated significant prediction bias at low sequencing coverages.

The average sequencing yield of 22.57 Gb represents a significant improvement on flow cell yield from previous ONT studies in cattle [26] and other species [27, 28]. This is largely a result of advances in our ability to fully utilise the sequencing capacity of each flow cell due to the release of a flow cell wash kit (Oxford Nanopore Technologies, Oxford). The wash kit is able to increase the yield from a single flow cell by unblocking ‘unavailable’ pores [29]. Pores on the flow cell can become blocked during sequencing by contaminants or tertiary DNA structures. To the best of our knowledge, the sequencing yield from one of our samples (41.13 Gb) is one of the largest MinION flow cell yields yet published. The proposed 2022 release of ONT’s shoebox sized PromethION P2 [30] could see portable sequencing flow cell yields increase 2-3-fold [31], through the increased density of pores on PromethION flow cells. This increase in the amount and speed of data generation would make *in-situ* genomic prediction more achievable, as the requirement to wait for data acquisition would be drastically reduced.

The average read length from these tail hair samples was significantly shorter than the average read length from previously sequenced tissues [26, 28]. The average Phred base quality of 20.52 represents a read error-rate of 0.9%, similar quality scores were reported by Runtuwene et al. [32] also using R9.4 flow cell chemistry. This shows the improvement in ONT sequencing from the initial reports using the R7.3 flow cell chemistry [32, 33] of base quality scores between 6.88 and 9.4 which relate to an error rate of 20.5% and 11.5% respectively. Further improvements are likely to arise with additional advances in base calling software, which have already shown significant progress [34, 35]. Continued improvements in base call accuracy will lead to more reliable genotypes, and therefore more accurate genomic predictions.

Although the sequencing effects of read length, base quality, effective mapping percentage and number of reads mapping with quality 0 had no significant effect on the ONT genomic prediction accuracy, this should still be investigated further. It is possible that a lack of variability between samples in this study made it difficult to model the true effect of these factors. Calculating genomic predictions using sequence data from different tissues or library preparation kits could help to investigate these different factors. It is particularly critical to test these effects under field conditions which reflect the real-world variation that may be experienced by either clinicians or producers applying this technology *in-situ*.

The non-imputed genotyping method performed significantly worse than either of the two imputation methods. This is likely because the missing genotypes were assigned as homozygous reference and therefore received no marker effect, because a homozygous reference is coded as a 0 in the genotype matrix, thereby giving 0 × the alternative allele effect. As the average sequencing coverage decreased the number of missing loci increased and therefore the breeding values were under-estimated at low coverages. This is supported by the decrease in the regression coefficient of ONT GEBV on SNP array GEBV for the non-imputed method as the sequencing coverage decreased. While not unexpected, this phenomenon has important implications: individuals with deeper sequencing coverage will have higher genomic prediction values, as more of their loci will be represented as non-zero in the genotype matrix. Therefor this method is unsuitable for any circumstance where the sequencing coverage is not tightly controlled.

The allele frequency method of imputation had the highest correlations with the SNP array breeding values across the range of coverages. However, overestimation of breeding values in animals at the lower end for each trait and under estimation of breeding values for animals at the high end was observed at low coverages (Figure 4). This type of prediction bias was also reported by Pimentel, Edel [36] who investigated the effect of imputation error on genomic predictions in 3,494 dairy cows genotyped on a low-density SNP array. They concluded this type of bias was an artifact of imputation algorithms suggesting the most frequent haplotype within a population whenever a haplotype cannot be observed unambiguously. A similar bias was likely seen here because assigning the missing genotypes using allele frequencies (i.e., the average genotype within the population) has regressed the breeding value of each animal toward the population mean. At high coverages (4x and 2x) this is less prominent because there are fewer missing genotypes. However, at the low coverages (1x and 0.5x) a significant number of markers are missing ONT genotypes and therefore the regression toward the mean for animals is more prominent. Therefore, this method is likely only useful when within-group rankings are of interest, and not absolute predictions: for example, if a producer wanted to identify the top 30% of animals for a certain trait within their herd.

The Beagle [25] imputation method for genotyping missing loci was the overall most accurate method when considering both correlation and prediction bias. Although the correlations for some traits were not as high as using the allele frequencies, the prediction bias was consistently close to 1 across all coverages and traits, indicating there was little over or under estimation of breeding values. This is likely because imputation with Beagle [25] uses linkage disequilibrium between genotyped markers to establish haplotypes. This means that the missing genotypes are imputed using the full extent of available information from the genotyped data (i.e., nearby markers genotyped with ONT reads). This approach is very different to imputing the missing genotypes using the allele frequency without considering nearby ONT genotyped markers. Imputation methods such as this are likely the most useful approach, despite the added time required to run the imputation program, which is currently substantially less time than required to produce the sequence data.

Here we tested the effect of a single imputation program, Beagle [25]. Further increases in the accuracy of ONT genomic predictions could be achieved using purpose built, low-coverage imputation packages such as the recently published program QUILT [37]. This particular imputation package produced similar genotype accuracies as a SNP array imputed using Beagle v.5.1 with 1x ONT sequencing coverage [37]. Additionally, imputation and genomic prediction accuracy could further be increased by genotyping genome wide SNPs rather than exclusively SNPs from the BovineHD BeadChip (Illumina, San Diego, CA) SNP array. This would have the advantage of incorporating the additional information available from whole genome sequencing (a very large number of polymorphic loci) rather than SNP array genotyping.

Other methods to increase the accuracy of the genomic prediction also include using sequence trimming quality control tools such as Prowler [38]. These tools increase the average Phred base quality of reads which reduces alignment errors and increases the number of effective reads. Studies based on short read information have found that optimisation of imputation strategies can have large effects on the accuracy of imputed genotypes [39, 40]. It is likely that appropriately optimised imputation strategies will increase the accuracy of imputation from ONT data further than what we have reported here and may allow for the calculation of accurate genomic predictions from ultra-low (less than 0.5X) sequence coverage, making the method more cost effective.

Targeted sequencing methods also hold enormous potential to increase the accuracy of genomic prediction using ONT sequencing. A promising method for *in-situ* applications is ONT’s adaptive sequencing, which can target loci within a genome for sequencing without molecular intervention [41, 42]. This method only requires the location of genetic markers and no additional laboratory steps, therefore would be ideal for on-farm or clinical applications of genomic prediction with the MinION [41, 42].

Although each flow cell was run for 96-hrs, the data used here for genomic prediction represent a small subset of the data acquired from each flow cell. For example, given the size of the bovine genome, 1x and 0.5x sequencing coverage represent 3 Gb and 1.5 Gb of sequence data respectively. This means that accurate genomic prediction for each animal would have been achievable in a fraction of the 96-hr sequencing run. Xu, Ge [43] reported an average yield of 0.48 Gb of sequence data per hour on a flow cell. For 0.5x sequencing coverage it would take a little over 3 hours to obtain sufficient data for accurate genomic prediction. With the above-mentioned optimisation steps, it is possible that the minimum required sequence data could fall further, decreasing the sequencing time. ONT also provide rapid library preparation protocols which are capable of preparing sequencing libraries in 15 minutes. This could make ONT genomic prediction far more rapid than traditional methods by eliminating the need to send samples to a laboratory. Studies have already demonstrated that ONT sequencing can significantly decrease turnaround times for pathogen identification by providing *in-situ* sequencing [44–46], genomic predictions could be the next step in *in-situ* genomic diagnosis.

Studies have demonstrated the ability for ONT data to characterise structural variants [47, 48], provide rapid pathogen identification [44–46] and assemble both large and small genomes [33, 35, 49]. In cattle, ONT data has successfully characterised the poll allele [26] and a novel structural variant in the *ASIP* gene controlling coat colour in Nellore cattle [50]. More recently in cattle, ONT sequencing was also used to annotate novel transcript isoforms by multiplex sequencing 32 bovine tissues on a single ONT flow cell [51]. Using cattle as an example agricultural species, we have successfully demonstrated yet another application of ONT data. Further optimisation of genotyping-by-sequencing methods with ONT sequence data by combining adaptive sequencing and optimised imputation methods, could see the required coverage for accurate genomic prediction decrease further. Given the average MinION flow cell yield achieved here, more than 20 human/cattle genome sized samples could be multiplexed on a single flow cell, making this approach cost-effective. This study is the first demonstration of genomic predictions using ONT sequencing, which has applications not only in agriculture, but in the clinical setting also.

## Materials & Methods

### UQ Ethics

Tail hair samples from 19 Droughtmaster animals were collected under the University of Queensland ethics approval numbers SVS/301/18 and SVS/465/18.

### DNA preparation

DNA for this study was extracted from tail hairs collected from 19 Droughtmaster heifers. The hairs were pulled and stored at room temperature for more than a year prior to DNA extraction. A subset of the hairs was used for genotyping on the 777k BovineHD BeadChip (Illumina, San Diego, CA). The remaining hairs were used for ONT sequencing. Genomic DNA for sequencing was extracted using the Gentra Puregene Tissue Kit (Qiagen) according to the manufacturer’s instructions with modifications. Briefly, 20-30 hair samples were lysed in 300 μl of Cell lysis solution (Gentra® Puregene® Tissue Kit) and 1.5 μl of Proteinase K solution (20mg/ml) for 5 hours at 55°C. RNA was then digested by addition of 1.5 μl of RNase A Solution, following 1-hour incubation at 37°C. Samples were placed on ice for 5 minutes after adding 100 μl Protein Precipitation Solution (Gentra® Puregene®Tissue Kit) and spun at 14000 x g for 3 minutes. 300 μl of Isopropanol was used to precipitate DNA. Samples were centrifuged at 14000 x g for 3 minutes. DNA pellets were washed in 300 μl of 70% ethanol, air-dried for 5 minutes and resuspended in 55 μl of DNA Hydration Solution (Gentra® Puregene® Tissue Kit).

DNA concentrations were measured using the Qubit dsDNA Broad Range assay kit (Thermo Fisher Scientific). The purity of the extracted DNA was determined with the NanoDropND 1000 (v.3.5.2, Thermo Fisher Scientific), assessing the 260/280 nm and 260/230 nm ratios. The size of extracted DNA was examined using pulsed-field gel electrophoresis (Sage science, USA) with a 0.75% Seakem Gold agarose gel (Lonza, USA) in 0.5X Tris/Borate/EDTA (TBE) running buffer, run for 16 hours at 75 V. The gel was stained after the electrophoresis with SYBR Safe dye (10000x) and visualized using Quantity One analysis software (Bio-rad).

### Sequencing methodology

Extracted DNA samples were prepared using a ligation kit (SQK-LSK 109, Oxford Nanopore Technologies) based on the manufacturer’s instruction with some modifications. Starting with 4 – 8 μg of DNA produced enough sequencing library to provide up to 4 flow cell loads from a single library. Samples were diluted at the clean-up points with nuclease-free water to prevent bead clumping during 0.4x AmPureXP purifications. End-prep reaction and ligation incubation times were increased to 30 minutes and 1 hour respectively. During the 96-hour sequencing run the flow cell was washed three times using the nuclease-flush kit (Oxford Nanopore Technologies) and then reloaded with the same library. The use of DNase in flushing and flow-cell re-fuelling helps to remove blocking DNA and increases the sequencing output.

### Genotyping methods

Base calls were made from the raw current disruption data using GUPPY (version 4.2.2, Oxford Nanopore Technologies) on the University of Queensland high performance computing infrastructure. The sequence data (fastq files) from each animal were randomly subsampled down to 4x, 2x, 1x & 0.5x sequencing coverage using a bash script. Minimap2 v2.17 [52] was used to align the reads to the *Bos taurus* reference genome ARS-UCDv1.2 [53], using the default ONT settings with the flag *-x ont-aln*. Samtools mpileup v1.3 [54] was used to create a pileup of the reads at 641,163 SNP loci that were also genotyped on the 777k BovineHD BeadChip (Illumina, San Diego, CA) and are segregating in Australia’s northern beef population.

A variable allele count genotyping method was used to genotype the SNP loci. Briefly, this method grouped loci with similar coverage together and called genotypes using a separate minimum allele count for each group (Figure 5). The minimum allele counts for each coverage group were chosen such that there was at least a 95% probability of observing both alleles given the random sampling nature of sequencing a diploid species.

**Figure 5:**
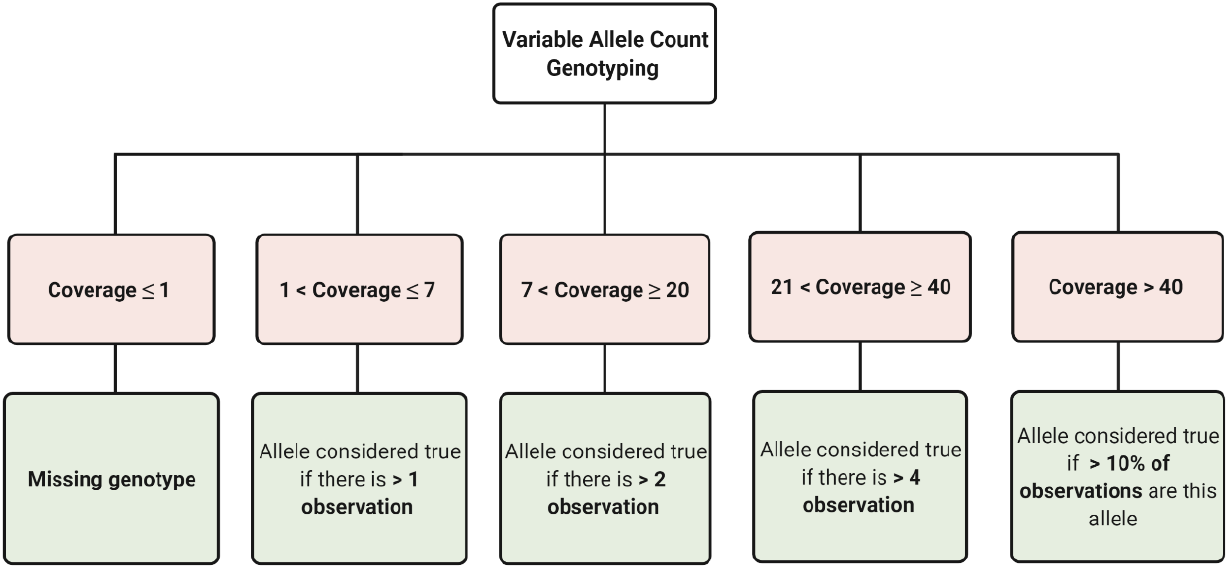
Variable allele count genotyping flow chart

Three methods were used to assign genotypes at loci with less than 2x coverage: non-imputed, imputed-AF and imputed-Beagle. The non-imputed approach assigned missing loci as homozygous reference, coded as 0 in the genotype matrix. The imputed-AF method used the allele frequency within the population, calculated from SNP array data, to genotype the missing loci. The sum of the probabilities of each genotype given Hardy-Weinberg equilibrium multiplied by the genotype codes, 0 for homozygous reference, 1 for heterozygous and 2 or homozygous alternate were used to fill missing genotypes. Using continuous genotype values between 0 and 2 for this method allowed some accounting for the uncertainty of genotyping from low coverage sequence data. Finally, the imputed-Beagle method used the imputation package Beagle v5.1 [25] to impute the missing genotypes given a reference panel of 1,200 animals which represented a subset of animals in the 1000 bull genomes project [54] with high *Bos indicus* content. The effective population and window parameters in Beagle were set to 100,000 and 100 respectively based on Pook, Mayer [56]. For imputation of the 0.5x sequencing coverage genotypes the window parameter was increased to 158 cM to ensure genotyped markers overlapped with the reference panel on each chromosome.

### SNP-BLUP method for EBVs

Genomic BLUP solutions from [24] were back solved to obtain SNP effects using GCTA [57] at 641,163 loci. The SNP effects were for four traits: hip height, body condition score, body weight and CL600 (presence of corpus luteum at 600 days). The presence of a corpus luteum is used as an indicator of heifer puberty in beef cattle [58, 59].

The estimated phenotypes 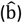 for the four traits were calculated using SNP array genotypes, contained in an *n* by *m* design matrix **M**, where *n* is number of animals and *m* is number of markers (641,163). The matrix contained 0, 1 or 2, designating homozygous, heterozygous and homozygous alternative genotype calls from the SNP array. The estimated phenotypes 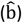 were then calculated using:

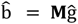

The ONT genotypes at the 641,163 loci were then used to calculate genomic breeding values for the four traits. The SNP marker information was contained in an *n* by *m* design matrix **N**, where *n* is number of animals genotyped using ONT data, and *m* is number of markers that were used (641k, corresponding to the same loci as in matrix **M**). The matrix contained genotype values between 0 and 2, according to the genotyping method used (non-imputed, imputed-AF or imputed-Beagle), and consistent with the nomenclature in matrix **M**. The matrix contained only 0, 1 or 2 for the non-imputed and imputed-Beagle methods, however contained continuous genotypes between 0 and 2 for loci imputed using the allele frequency. The estimated phenotypes 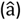 were then calculated using the same estimated marker effects as were used for the SNP array data:

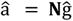

Linear models were used to evaluate the correlation between 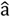 and 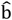, being the estimated genetic value derived from ONT and SNP array, respectively. The correlation between 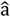 and 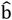 was reported as well as the prediction bias (regression coefficient of the SNP array-GEBV on the ONT-GEBV).

Linear models were also used to evaluate the effect of read length, base quality, effective mapping percentage and number of reads mapping with Phred mapping quality 0 on ONT genomic prediction accuracy. To evaluate each covariate, **y** was calculated as the residuals from the model 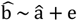, where 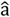 was an array of SNP-array based genomic predictions (from above), and 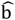 was an array of ONT-based genomics predictions (from above). The significance of each covariate (read length, base quality, effective mapping percentage or number of reads mapping with Phred mapping quality 0) on the genomic prediction accuracy from ONT sequence data was modelled using:

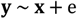

where **x** was an array of the covariate values (read length, base quality, effective mapping percentage or number of reads mapping with Phred mapping quality 0) as a random effect; and **y** was the difference between the SNP-based genomic prediction and the ONT-based genomic prediction.

## Acknowledgements

The authors would like to acknowledge the many contributions from friends and colleagues. Firstly, we would like to thank the 54 collaborators in the Northern genomics project, as well as Mr James Copley, Ms Shannon Speight and Dr Geoffrey Fordyce, who were instrumental in its coordination. We would also like to thank Professor Michael McGowan and Dr Russell Lyons for their help in collecting the tail hair samples for sequencing. We are grateful for funding from the MLA Donor Company and University of Queensland through projects P.PSH.P0833, B.STU.2001, L.GEN.1713 and L.GEN.1808.

## Notes

### Competing Interest Statement

The authors have declared no competing interest.

